# Development and Application of Multidimensional Lipid Libraries to Investigate Lipidomic Dysregulation Related to Smoke Inhalation Injury Severity

**DOI:** 10.1101/2021.10.13.464246

**Authors:** Kaylie I. Kirkwood, Michael W. Christopher, Jefferey L. Burgess, Sally R. Littau, Brian S. Pratt, Nicholas Shulman, Kaipo Tamura, Michael J. MacCoss, Brendan X. MacLean, Erin S. Baker

## Abstract

Lipids play many biological roles including membrane formation, protection, insulation, energy storage, and cell division. These functions have brought great interest to lipidomic studies for understanding their dysregulation in toxic exposure, inflammation, and diseases. However, lipids have shown to be analytically challenging due to their highly isomeric nature and vast concentration ranges in biological matrices. Therefore, powerful multidimensional techniques such as those integrating liquid chromatography, ion mobility spectrometry, collision induced dissociation, and mass spectrometry (LC-IMS-CID-MS) have recently been implemented to separate lipid isomers as well as provide structural information and increased feature identification confidence. These multidimensional datasets are however extremely large and highly complex, resulting in challenges in data processing and annotation. Here, we have overcome these challenges by developing sample-specific multidimensional libraries using the freely available software Skyline. Specifically, the human plasma library developed for this work contains over 500 unique, experimentally validated lipids, which is combined with adapted Skyline functions for highly confident lipid annotations such as indexed retention time (iRT) for retention time prediction and IMS drift time filtering for increased sensitivity and selectivity. For broad comparison with other lipidomic studies, this human plasma database was initially used to annotate LC-IMS-CID-MS data from a NIST SRM 1950 extract, giving comparable results to previous studies. This workflow was then utilized to assess matched plasma and bronchoalveolar lavage fluid (BALF) samples from patients with varying degrees of smoke inhalation injury to identify potential lipid-based patient prognostic and diagnostic markers.

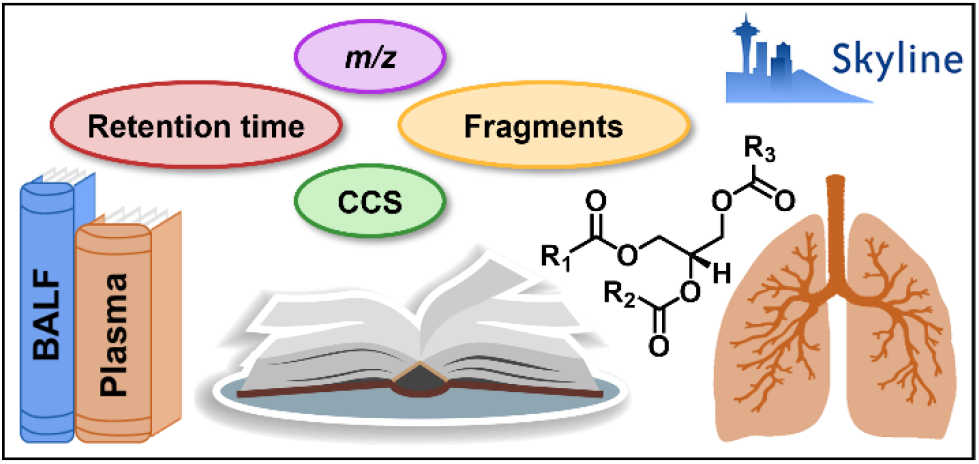

## INTRODUCTION

While lipids are an essential component of the cellular membrane, they are also an extremely diverse class of biomolecules with roles extending beyond just cellular signaling, membrane structure, and energy storage.^1, 2^ Lipid dysregulation has therefore been linked to diseases such as cancer, cardiovascular disease, diabetes, and other metabolic disorders as well as toxic exposures and environmental perturbations.^3–5^ Thus, recent studies have focused on understanding lipid functions and biological mechanisms for their use as biomarkers or to create novel disease treatments and thoroughly evaluate system perturbations. However, lipidomic studies are often challenged by the extensive concentration range lipids exhibit in biological samples as well as the separation and data annotation challenges associated with the structural similarity and isomeric nature of lipids.^6^ For example, thousands of lipid isomers exist which have the same elemental composition but differ in structural arrangements such as variation in head group and alkyl chain makeup, stereochemistry (*sn*-position), or double bond position and orientation.^7–9^ These similarities have driven improvements to analytical strategies for lipidomic studies over the last decade. To achieve more comprehensive lipidomic characterization, powerful analytical tools beyond traditional LC-MS-based methods have been employed including the use of multidimensional separation techniques such as LC-IMS-CID-MS. IMS is a gas-phase separation technique in which ions are separated by size, shape, and charge. While IMS has many different types (e.g., traveling wave IMS, trapped IMS, field asymmetric IMS, etc.) in this work drift tube IMS (DTIMS) was specifically used and will only be discussed from here on. In DTIMS, a drift cell is filled with a buffer gas such as nitrogen which collides with ions as they traverse the tube under an applied electric field. This causes smaller ions to have fewer collisions and migrate faster that more extended ions, thus separating the sizes by their drift time. An ion’s drift time can then be correlated to its gas-phase ion-neutral cross sectional area (CCS).^10^ These IMS evaluations therefore allow for isomer distinction, separation of biomolecular classes and background interferences due to backbone differences, and increased annotation confidence via an additional parameter for feature matching.^7, 8, 11^ Additionally, since DTIMS is performed in the gas-phase, its millisecond separations are easily performed sandwiched between frontend chromatography systems and following time-of-flight mass spectrometry analyses for multidimensional assessments of each detected ion.^12^

While multidimensional separations offer valuable benefits, the resulting datasets can be quite difficult to annotate, particularly for complex sample types such as biofluids or tissues. Thus, users are often faced with the challenge of manually annotating their data, which can only be done for a limited number of targets in small datasets or using rapid processing software that may disregard the IMS dimension altogether. The freely available and open-source software Skyline has traditionally been used for LC-MS-based proteomics data processing, however it was recently adapted to process IMS data from multiple instrument vendors for both proteomics and small molecule data.^13, 14^ We therefore leveraged these advancements as well as additional Skyline adaptations to develop multidimensional lipid libraries for human plasma and BALF analyses. Here we make the lipid database publicly available including an editable spectral library storing LC-IMS-CID-MS reference information for 516 unique lipids and an indexed retention time (iRT) prediction calculator to account for retention time shifts.^15^ This workflow was then applied to human plasma and BALF analyses to annotate lipidomic LC-IMS-CID-MS data and evaluate the information from each biofluid and the lipid dysregulation related to smoke inhalation severity. Since smoke inhalation causes injury and malfunction of the respiratory tract, it is one of the strongest determinants of burn patient mortality beyond age and burn size.^16^ Inhalation injury also increases morbidities in burn patients and is the source of over 23,000 injuries annually.^17^ In addition to direct thermal and heat damage, smoke consists of highly toxic combustion products such as carbon monoxide, cyanide, phosgene, formaldehyde, hydrogen sulfide, ammonia, and up to 400 others (**Figure 1**).^18^ These stressors can each contribute to a range of outcomes including swelling, cytokine accumulation and other inflammatory responses, increased alveolar permeability, acute pulmonary edema, acute lung injury, and acute respiratory distress syndrome (ARDS), among others (**Figure 1**).^16^ However, the molecular response to smoke inhalation is not well understood, and identifying prognostic and diagnostic factors in patients with inhalation injury has proven to be challenging.^19–21^ In this work, we characterized lipid dysregulation for both the matched plasma and BALF from smoke inhalation patients with varying levels of severity. This allowed us to gain insight into each sample type, assess the molecular mechanisms of injury and potentially identify lipid markers to enhance clinical estimations of prognosis and enable better treatment diagnosis.

**Figure 1.**
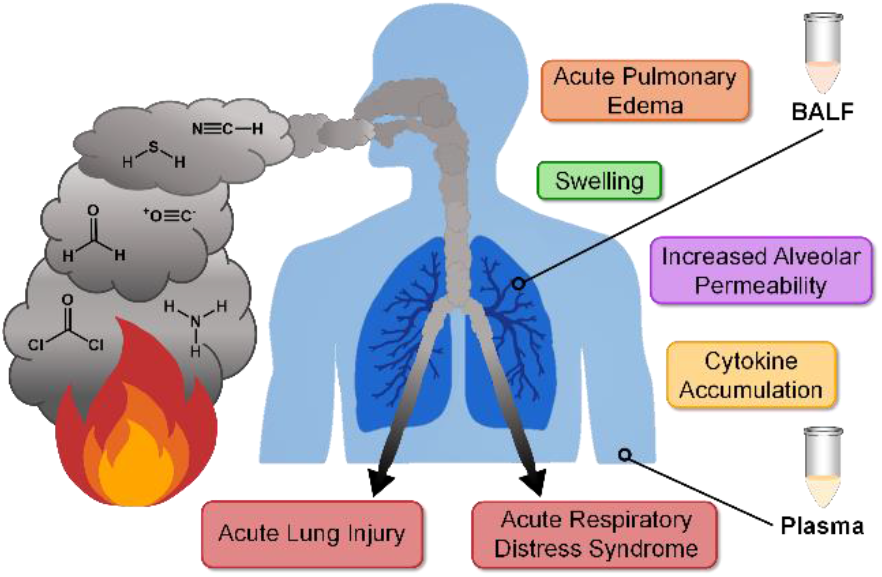
Overview of select toxic components of smoke, inhalation injury outcomes, and the sample types studied in this work.

## EXPERIMENTAL METHODS

### Sample Collection and Preparation

Matched plasma and BALF samples were obtained from 21 adult subjects sustaining smoke inhalation injuries. Of the 21 patients, 6 unfortunately died from their injuries. All patients were admitted to a burn center and intubated within 24 hours of admission. All samples were deidentified and informed consent was obtained from all participants. A bronchoscopy was performed within 6-12 hours post-injury to collect BALF specimens. The BALF samples were then dithiothreitol (DTT)-treated and protein concentrations were determined by measuring the absorbance at 280 nm. All BALF samples were then stored at −80°C until LC-IMS-CID-MS analysis. Blood samples were simultaneously collected and processed with the BALF, then spiked with EDTA and stored at −80°C until analysis. Lipids were then extracted from both sample types using a modified Folch extraction.^22^ First, 50 μL of EDTA plasma or the volume needed to reach a protein amount of 250 μg for DTT-treated BALF was mixed with 600 μL of 2:1 −20°C chloroform/LC-MS grade methanol. Samples were vortexed for 30 s, then phase separation was induced by adding 150 μL of LC-MS grade water and vortexing for an additional 30 s. Samples were left on ice for 5 min, then centrifuged at 12,000xg and 4°C for 10 min. 300 μL of the bottom lipid layer was then transferred and solvent was evaporated using a SpeedVac. Extracts were reconstituted in 10 μL of chloroform then mixed with 190 μL of LC-MS grade methanol prior to analysis. Pooled quality control (QC) samples made from the equal same volumes of each sample were also extracted alongside the experimental samples. The NIST SRM 1950 (metabolites in frozen human plasma) was extracted in triplicate in the same fashion as the experimental EDTA plasma samples.

### LC-IMS-CID-MS Analyses

Lipid extracts were analyzed using an Agilent 1290 UPLC system coupled with an Agilent 6560 IM-QTOF platform (Santa Clara, CA) with the commercial gas kit and an MKS Instruments precision flow controller (Andover, MA). Both positive and negative mode ESI data was collected with a mass range of *m/z* 50-1700. Lipids were first separated by reversed-phase liquid chromatography by injecting 10 μL onto a Waters (Milford, MA) CSH column (3.0 mm x 150 mm x 1.7 μm particle size). The 34-minute gradient having a flow rate of 250 μL/min and mobile phase A of 10 mM ammonium acetate in 40:60 LC-MS grade acetonitrile/water and mobile phase B of 10 mM ammonium acetate in 90:10 LC-MS grade isopropanol/acetonitrile is detailed in **Table S1**. Lipid ions were fragmented by collision-induced dissociation using an all ions data-independent acquisition approach, where the collision energy was ramped based on the precursor ion’s drift time (**Table S2**).^23^ Brain total lipid extract (BTLE) from Avanti Polar Lipids (Alabaster, AL) was used as an additional QC sample to monitor instrument performance.

### Library Development

Lipid libraries were developed from pooled QC plasma and BALF samples in both positive and negative mode within Skyline software (Skyline-daily 20.2.1.315 through 20.2.1.414). Plasma spectral libraries were exported as .blib files. The libraries were populated with lipid class, name, precursor formula, and adduct(s); product and neutral loss formulas, adducts, and relative abundances; and CCS and high energy drift offset values. Lipid shorthand nomenclature followed the guidelines proposed by the LIPID MAPS consortium.^24^ Lipid transition lists were generated manually or using LipidCreator, and the observed fragmentation patterns were validated via comparisons to literature and in-silico spectra when available.^25–27^ All lipids included in the libraries were manually validated and met the confidence criteria of presence in more than one sample, ±5 ppm mass measurement accuracy, co-elution of multiple adducts (if applicable), retention time within the expected class-specific or reported window, drift alignment of precursor and fragment ions, and CCS values within 2% of database values (if present).^15^ iRT calculators were then built from the positive and negative mode pooled plasma data using sets of 20 lipids spanning the LC gradient as the endogenous reference lipids. All plasma library lipids were assigned iRT values on a 0-100 scale using a Lowess regression.^15^

### Data Processing and Statistical Analysis

Data files were single-field calibrated using the Agilent ESI tune mix in Agilent IM-MS Browser software.^10^ The calibrated data was annotated using Skyline software (Skyline-daily 20.2.1.315 through 21.1.1.223) with the sample-specific libraries for the target lists. Retention times for each library lipid in the experimental samples were predicted within a 2-minute window using the developed iRT calculators. Drift time filtering was used with a resolving power of 50. All annotations were manually validated. For lipids that were detected as multiple adducts and/or in both ionization modes, one representative target (i.e., adduct) was chosen for statistical analysis. The extracted ion intensities were exported to Excel for further analysis.

Metaboanalyst was used to normalize exported lipid abundances to the sample total ion current (TIC) value and apply a log10 transformation.^28^ Statistical analysis was performed using pmartR.^29^ First, outliers were determined and removed based on an RMD-PAV algorithm, Pearson Correlation, and PCA. Subject age and sex were assigned as covariates. Smoke inhalation injury subjects were grouped by survival or mortality to compare lipid abundances and identify statistically significant changes between the groups using an ANOVA with a Holm multiple comparison correction and significance cutoff of p < 0.05.

## RESULTS AND DISCUSSION

### Lipid Library Composition

Multidimensional lipid libraries were built from LC-IMS-CID-MS data of pooled human plasma and BALF lipid extracts using Skyline. Each library lipid was manually validated with the criteria of 1) presence in multiple samples, 2) high mass measurement accuracy (±5 ppm), 3) co-elution of multiple adducts (if applicable), 4) retention time within the class-specific window or matching literature values, 5) presence of IMS drift-aligned fragment ions, and 6) calculated CCS values within 2% of previous database values (if known).^15^ The human plasma lipid library contains over 6100 transitions for 854 lipid targets corresponding to 516 unique lipids as some were observed as different ions in both positive and negative ion mode (**Figure 2**). The human BALF library contained over 3400 transitions for 460 targets, corresponding to 377 unique lipids. A high degree of overlap was observed between both sample types, resulting in a total of 542 unique lipids and 890 lipid targets, of which approximately 93% are not reported in the LIPID MAPS Structural Database via the CCS Compendium.^11^ These 542 lipids span 18 classes from 5 of the 8 lipid categories as defined by LIPID MAPS including fatty acyls (AC, ANA, FA, FAHFA) glycerolipids (DG, TG), sphingolipids (Cer, GM3, HexCer, SM), sterol lipids (CE), and glycerophospholipids (PA, PC, PE, PG, PI, LPC, LPE) including plasmalogen species.^30^ Additionally, the alkyl chains range from 10-26 carbons and 0-6 degrees of unsaturation. The lipid targets also include protonated and deprotonated species as well as sodium-, ammonium-, acetate-, and formate-adducted species.

**Figure 2.**
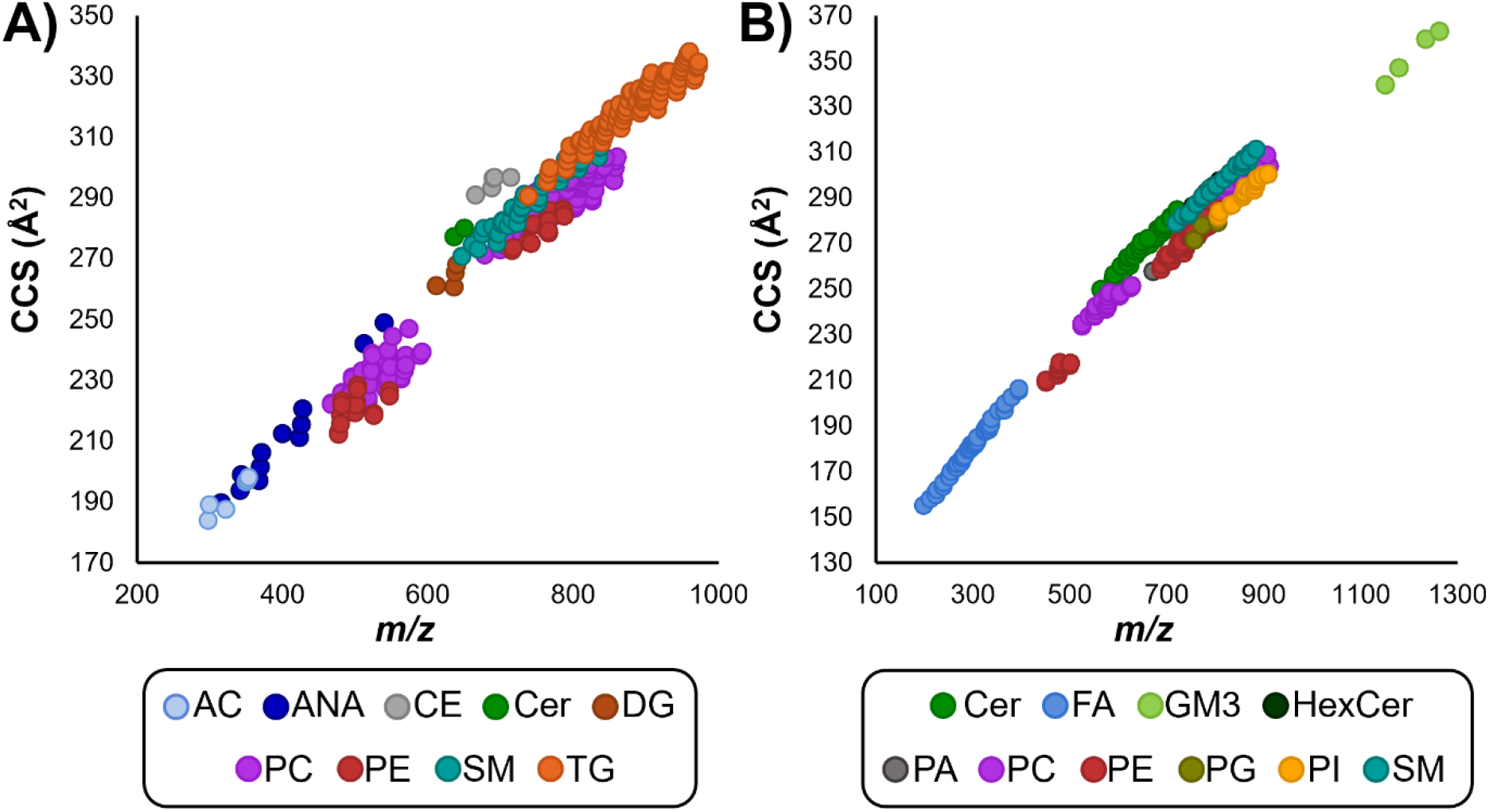
Conformational space of the plasma lipids included in the **A**) positive and **B**) negative ionization mode libraries. All 854 lipid features, corresponding to 516 unique lipids when accounting for multiple ion types, are plotted by CCS versus *m/z* value. These lipids span 18 classes from 5 lipid categories. Features are colored by class to emphasize class-specific trendlines.

In all cases, the number of unique lipids is lower than the number of library targets due to many lipids being detected as multiple ion types in both ionization modes. Additionally in some cases there were multiple features that could not be distinguished, possibly resulting from *sn* connectivity or double bond differences, which were therefore assigned to a single lipid and denoted with a or b following the lipid name, based on the relative retention time or CCS order. On the other hand, there were some features where the fatty acyl chains could not be assigned due to low fragmentation efficiency or where multiple lipid isomers co-eluted. In these cases, the features were respectively annotated with the sum composition (i.e., total number of fatty acyl carbons and double bonds) or as multiple lipids separated by a semicolon. As plasma samples are more easily obtained, commonly evaluated, and more lipid rich than BALF, the plasma data was utilized to build a comprehensive, publicly available, and editable database comprised of a Skyline spectral library and iRT calculator. Therefore, both libraries (plasma and BALF) contain information on lipid class, name, precursor formula, and adduct(s); product and neutral loss formulas and adducts; and CCS and high energy drift offset values. The human plasma lipid database also contains reference CID spectra in the form of a Skyline spectral library and normalized retention time (iRT) values for retention time prediction which will be discussed in detail below. **Figure 2** displays the lipid composition of the human plasma library in the form of CCS versus *m/z* plots for both the positive and negative mode ions. These plots demonstrate both the diversity of lipid coverage as well as the ability of IMS-MS to separate and correlate lipids by class. Previous studies have quantitatively characterized the structural trends observed here.^7, 31, 32^ Briefly, conformational ordering is based on the relative size per mass unit of the molecular class of interest. Lipids are much larger in size than ring-based metabolites of similar mass due to the long alkyl chains, thus they occupy a unique space within the IMS dimension. Similarly, each lipid category and many lipid classes can be distinguished using this approach. Lipid classes give unique linear trendlines which can be utilized for increased identification confidence and rapidly classifying unidentified lipid features. Additional information can be ascertained by further examination of each lipid class trendline. For example, the PC and TG trendlines in **Figure 2A** show sub-trends where the CCS of lipids of the same carbon chain length increase linearly with higher degrees of unsaturation. This phenomenon is discussed at length by Leaptrot and coworkers.^7^

### Lipid Database Evaluation

To evaluate the performance of our Skyline lipid database using a commonly analyzed sample, the National Institute of Standards and Technology (NIST) Standard Reference Material (SRM) 1950 – Metabolites in Frozen Human Plasma was extracted and analyzed in triplicate with the same LC-IMS-CID-MS platform used to create the database. The resulting data was then annotated in Skyline using the plasma lipid database to compare the detected lipids to previous studies that have deeply characterized SRM 1950. In one previous study, the LIPID MAPS consortium identified a total of 588 lipids using multiple extraction and analysis methods.^33^ A subsequent interlaboratory study by Bowden et al. identified 399 lipids by at least 5 of the 31 participating laboratories.^34^ Using the workflow developed for this study, we identified 483 unique lipids, a highly comparable number to both previous studies. This also corresponded to 94% of the total number of lipids included in our plasma spectral library. While the LIPID MAPS and interlaboratory studies reported lipids as the sum composition level, many of our unique identifications arise from the inclusion of lipid isomers. Both prior studies employed a wide variety of lipid category- and class-specific extractions and separation techniques including RPLC, HILIC, and GC, while our study was just with one extraction and an RPLC front-end. Furthermore, the composition of lipids identified here in the SRM 1950 by category is comparable to the previous findings, with a preference for categories that are well-extracted and separated with the method used here as opposed to those that may require specialized extractions, derivatization, or GC separations such as those for sterols (**Figure S1**). Thus, the plasma lipid database developed here is a powerful tool when coupled with an LC-IMS-CID-MS platform, providing hundreds of confident annotations using a single sample extraction and analysis platform.

### Skyline Functions Leveraged for the Lipid Database

For this study, we are also releasing the human plasma database including the Skyline spectral library and iRT calculator at https://panoramaweb.org/baker-lipid-ims.url to the scientific community. This library and its associated workflow leverages many new aspects of the Skyline small molecule interface to provide rapid multidimensional data processing and confident lipid annotations.^13^ One such capability is to build and view small molecule spectral libraries. Spectral libraries were created for plasma lipidomic data in both ionization modes to provide reference CID spectra when annotating experimental data. In the spectral library view, users can browse the CID spectra with annotated fragment ion signals (**Figure S2**), in addition to viewing the target lipid name, formula, adduct, *m/z* and CCS values, and observed retention time.

The Skyline iRT tool was recently adapted to standardize and predict small molecule LC retention times to account for run-to-run LC shifts, system differences, and method gradient time changes.^15^ In each ionization mode, a set of 20 endogenous references lipids seen in many different sample types and that spanned the LC gradient were assigned iRT values between 0-100 to calibrate the iRT calculator (**Table S3**). All remaining lipids in the library were subsequently assigned iRT values by Skyline based on their retention times relative to the reference lipids. When applied to experimental data, the user verifies the retention times of the 20 reference lipids in each sample, then the retention times of the hundreds of remaining lipid targets are predicted. Previous predictions using iRT for proteomic data in Skyline have been based on linear regressions, but due to the varying relative shifts in retention times when gradient times are shortened, in our lipidomics study a non-parametric locally weighted scatterplot smoothing (Lowess) model was applied to predict analyte retention times more accurately. **Figure 3** demonstrates the predictive capabilities of the developed plasma iRT calculator. Here, the iRT calculator was created from the pooled human plasma QC samples and applied to the experimental samples using the same 38-minute LC gradient method. To evaluate the predictions for the same gradient but different columns, the calculator was applied to commercially available brain total lipid extract (BTLE) data collected with a different LC column and instrument (and even at a different laboratory) but using the same mobile phase composition and LC gradient method. The observed predictions were highly accurate with an average of 2% difference, or within 0.6 minutes, from the observed retention times across the three replicates averaged and shown as a single dot in **Figure 3A**. When the run time was halved to 14 minutes, the predictions gave an average of 3% difference, or were within 1.1 minutes, from the observed retention times across three replicates (**Figure 3B**). Further training could be done at similar short gradients, but this fitting allows rapid comparison if the gradient times are drastically changed. **Figure 3C and 3D** demonstrate the utility of iRT for isomer separations, where the *sn*-1 and *sn*-2 isomers of LPC 18:1, which differ only by their connectivity to the stereocenter of the glycerol backbone, are separated by LC. These isomers have identical fragments and only small size difference given the resolving power of this IMS system (CCS values <2% difference), thus their LC separation is helpful but can lead to challenges with correctly annotating the two features. Based on literature retention times, the first eluting peak was annotated as LPC(0:0/18:1) (**Figure 3C**) and the second as LPC(18:1/0:0) (**Figure 3D**).^8^ Users of the plasma iRT calculator can therefore confidently annotate these features without the need of chemical standards or extensive literature searches using the iRT predicted retention times. While the iRT calculator was optimized for plasma lipidomic evaluations, it can be utilized for other samples as long as a majority of the 20 reference lipids are present. Additionally, labeled standards like those present in the Avanti Polar Lipids SPLASH LIPIDOMIX can also be used for this task.

**Figure 3.**
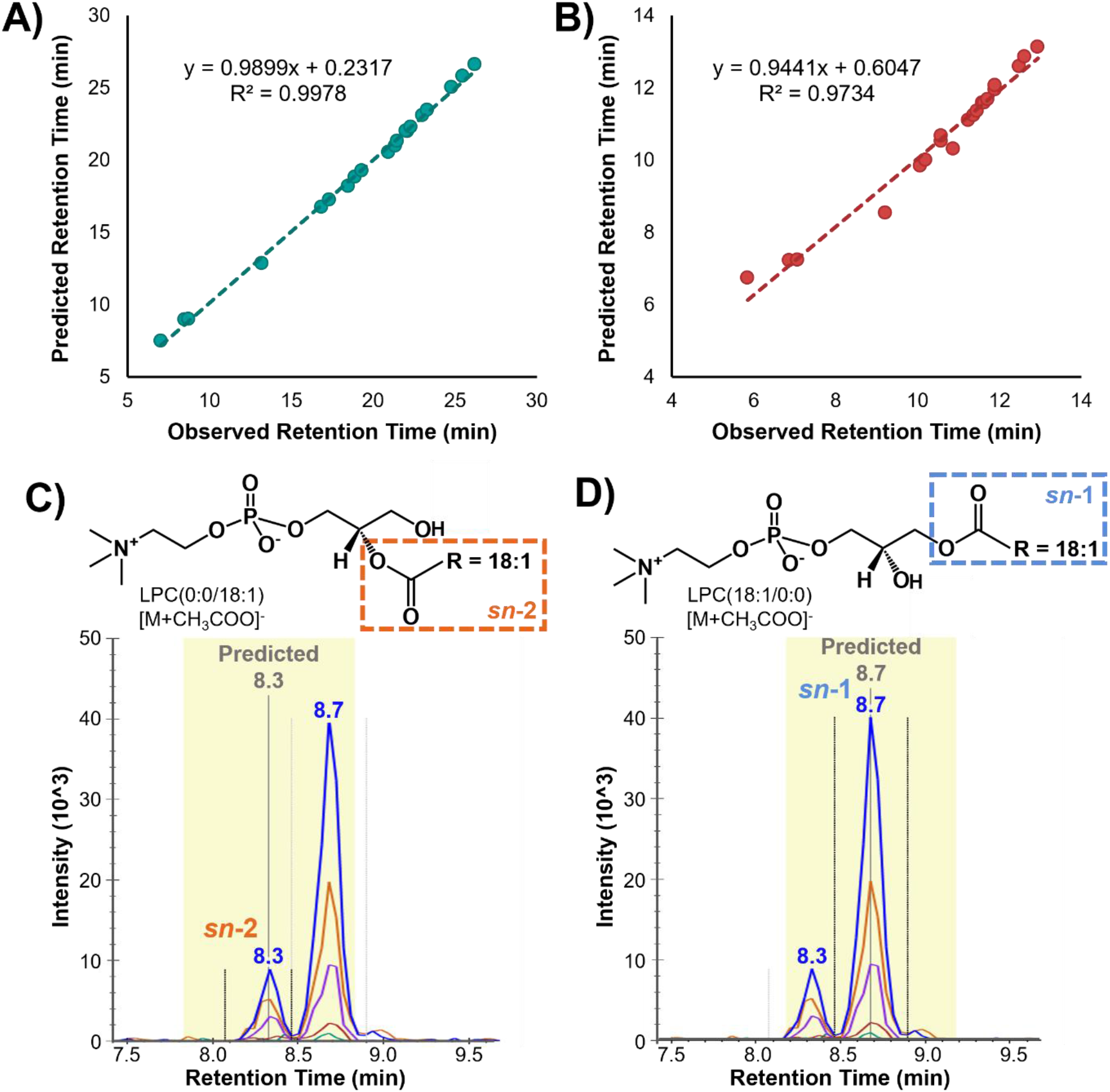
Predictive capabilities of the plasma iRT calculator. Accuracy of the iRT retention time predictions plotted as predicted versus observed retention times for triplicate BTLE data collected on a different column and instrument with the same mobile phase composition and LC gradient method at **A**) the original run time of 38 minutes and **B**) a shortened run time of 14 minutes. iRT prediction utility in assigning LC-separated isomers such as **C**) LPC(0:0/18:1) and **D**) LPC(18:1/0:0) as shown by the predicted grey bar.

A traditional capability of Skyline is the visualization and parallel analysis of multiple adducts for a single molecule.^13^ This function is leveraged here to increase the confidence of lipid annotations via the co-elution of multiple unique adducts for a given lipid class (**Figure S3**). For example, ceramides produce deprotonated as well as formate- and acetate-adducted species in negative ionization mode which can be utilized in their annotation, even though only one target is needed for relative quantitation.^26^ Annotation of nontraditional adducts can also increase the level of speciation achieved for a given lipid. For example, in positive mode, PC lipids form highly abundant [M+H]^+^ ions, but CID produces very low abundance neutral loss fragments which leads to many PCs being reported at the sum composition level instead of assigning the fatty acyl components.^26^ Sodiated PC ions are often disregarded as they are generally at least an order of magnitude lower in abundance than the protonated form. However, sodiated PCs give more abundant diagnostic fragments when using CID.^35, 36^ Here we found that at least 10% of the total abundance (precursor and fragment peak areas) of sodiated PCs was fragment ions that could be used to elucidate the fatty acyl composition compared to less than 0.25% of the total protonated PC abundance (**Figure 4**). The sodiated targets can therefore be utilized to assign fatty acyls and achieve higher speciation for the observed PCs, while the co-eluting deprotonated species can be used for downstream analyses.

**Figure 4.**
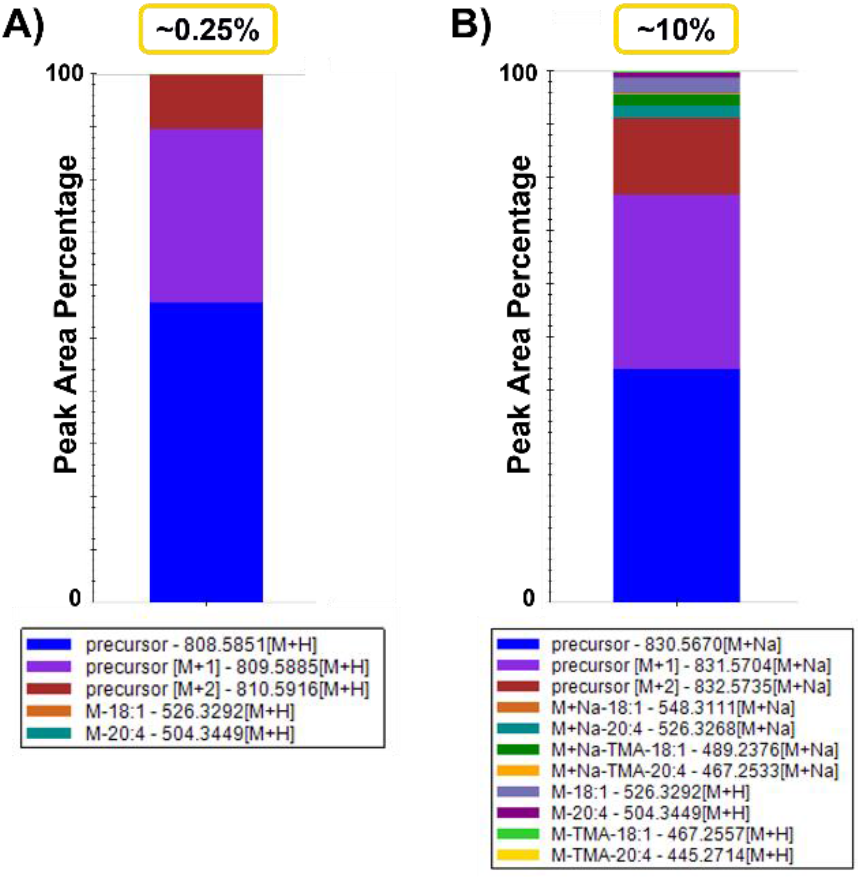
Use of sodiated phosphatidylcholines for fatty acyl assignment of PC(18:1_20:4). Example of peak area percentages for **A**) protonated and **B**) sodiated PC species in positive mode.

The IMS CCS value for each lipid in the library also provides many capabilities for their evaluation and confident identification. Each lipid target possesses a stored precursor CCS value which is used in conjunction with the single-field calibration parameters to back-calculate ion drift times.^10^ These drift times can then be used for drift time filtering, where Skyline generates a window of drift times in which signal will be extracted based on the IMS instrument resolving power. This function is used to increase sensitivity by filtering noise and interferences and when demonstrated by MacLean and coworkers, resulted in BSA peptide calibration curves with both lower detection limits and greater linearity.^14^ This drift time filtering of lipid signals is particularly useful in the removal of interferences due to commonly detected isomers and isobars which can contribute to a lipid of interest’s extracted ion intensity and alter subsequent lipid abundance comparisons. While MS1 drift time filtering is useful, this function is especially advantageous for filtering tandem MS data, or in this study data-independent acquisition (all ions) CID data, to annotate lipid features more accurately. The assignment of fatty acyl chains with DIA is a difficult task without drift time filtering, as diagnostic fragment ions are shared across most lipid classes. For example, lipids with an 18:0 fatty acyl chain will give the same diagnostic 18:0 transitions, particularly in negative mode where FA 18:0 [M-H]^-^ fragments are easily observed.^26^ Furthermore, highly abundant lipid fragment signals can dominate the majority of the chromatogram as shown in **Figure 5A**, where the teal and orange chromatogram traces are common FA 18:0 and 20:4 [M-H]^-^ fragments. Using drift time filtering significantly reduces these signals (**Figure 5B**), rendering the PE fatty acyl assignment more straightforward. Drift time filtering also improved the accuracy of the spectral library fragment ion relative abundances, as the library was generated from experimental data rather than neat standards with fragments arising from a single lipid. Skyline can also store a high energy drift time offset value within the spectral library entry for each lipid. This value accounts for the slightly lower drift time fragment ions result at compared to the precursors due to the slightly higher velocity imparted into the fragment ions from the voltage applied in the CID cell. Skyline is therefore able to shift the drift time filtering window, in this case by −0.5 ms, to account for the fragment ions arriving at the detector slightly faster than the larger precursor ions. The high energy drift spectra for the FA 18:0 [M-H]^-^ fragment at 283 *m/z* is shown in **Figure 5C** where all signal outside of the purple horizontal box, or aside from approximately 34.4-36 ms, is filtered out. Without drift time filtering, at least two visible ions would contribute to the extracted intensity of this fragment. Additional complexity arises from isotopic overlap of fatty acyl fragments of differing degrees of unsaturation. In the case of **Figure 5C**, if there were an abundant co-eluting lipid with a FA 18:1 [M-H]^-^ fragment (281 *m/z*), the [M+2] signal could interfere with the 18:0 signal.

**Figure 5.**
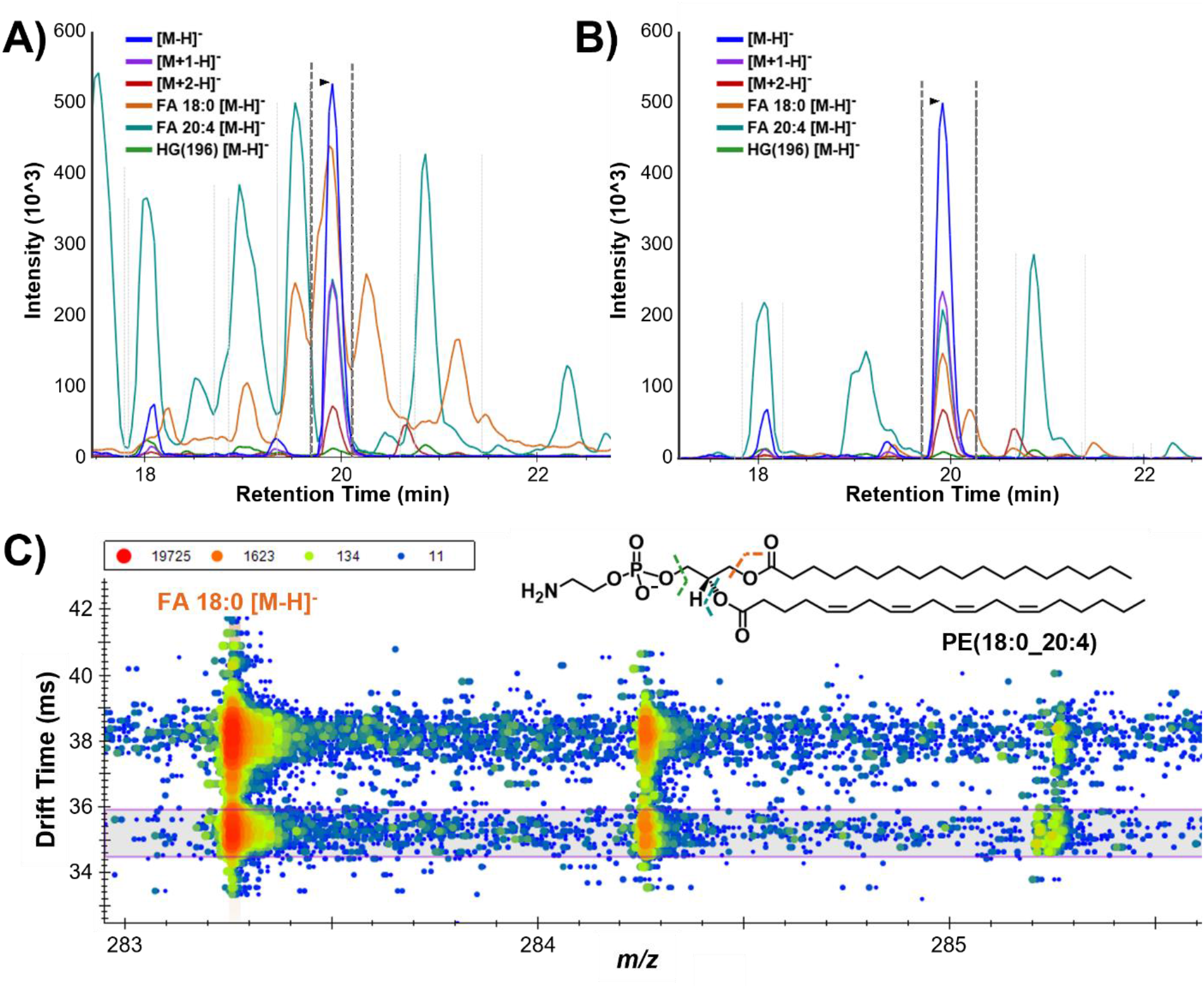
Drift time filtering for target lipid features in Skyline. Extracted ion chromatogram for the PE(18:0_20:4) precursor, M+1 and M+2 isotopes, and three characteristic fragments illustrate how **A**) before drift time filtering there are numerous and **B**) following drift time filtering a majority of the peaks are removed since they are not affiliated with the target ion. **C**) Fragment DIA drift spectra for FA 18:0 [M-H]’, where the signal outside of the purple drift time window belongs to another precursor ion and is filtered out, thereby decreasing interference and improving quantitative capabilities.

When comparing Skyline with other free, vendor-neutral lipid data annotation software tools, we noticed a majority do not support the IMS dimension.^37–39^ In cases where IMS information is utilized, lipids are often annotated prior to matching the observed precursor CCS value to either literature values, *in silico* predictions, or class-specific trendlines. Therefore, even multidimensional data may be subject to the incorrect assignment of fatty acyl chains due to co-eluting precursors if the software does not filter by CCS or check for drift alignment of product ions prior to annotation. Since the drift time filtering function can be utilized for traditional precursor and fragment ions as well as neutral loss fragments, this is an extremely important added functionality in the Skyline small molecule interface for lipidomic analyses as neutral losses are common for some lipid classes.^26^

### Lipid Markers of Smoke Inhalation Injury Severity

To apply our Skyline lipidomic libraries to real samples, multidimensional assessments were carried out on an LC-IMS-CID-MS platform for both subject-matched plasma and BALF samples from patients with smoke inhalation injury. The main goals of this study were to evaluate which sample type was best to identify potential markers of severe smoke inhalation injury severity and determine if specific lipid biomarkers could be found for subjects who did or did not survive their injury while controlling for age and sex. After each sample was analytically assessed, the data were analyzed using the plasma and BALF Skyline libraries discussed above which yielded 424 unique lipid identifications across five lipid categories in the samples studied. This corresponds to 78% of the lipids from the total combined libraries, as some were present in the pooled QCs used to build the libraries but were unable to be confidently identified in experimental samples, particularly in the BALF samples which showed a higher degree of variability.

A total of 25 lipids were found to be statistically significant between the two subject outcomes with 19 in plasma and 6 in BALF (**Figure 6A**). Since 3x more statistically significant lipids were observed in the plasma samples, this indicates it is a better biofluid for prediction of mortality. Additionally, we observed some blood in the BALF samples and since extensive airway injury can occur after smoke inhalation, added variability was noted in the samples as compared to plasma. Of the lipids observed as statistically significant in BALF, 2 were upregulated (FA 14:0 and AC 14:0) and 4 were downregulated (FA 25:0, PE(16:0_18:0), PC(15:0_16:0), PC14:0_16:0)) in subjects whose injuries led to mortality (**Table S4**). Specifically, dysregulation was observed among BALF FAs and PCs as well as an AC and a PE. A noteworthy observation was that while FA 14:0 and AC 14:0 were upregulated a PC containing a 14:0 fatty acyl chain was downregulated, indicating an increase in free FA (FFA) 14:0 in BALF from subjects with the most severe injuries.

**Figure 6.**
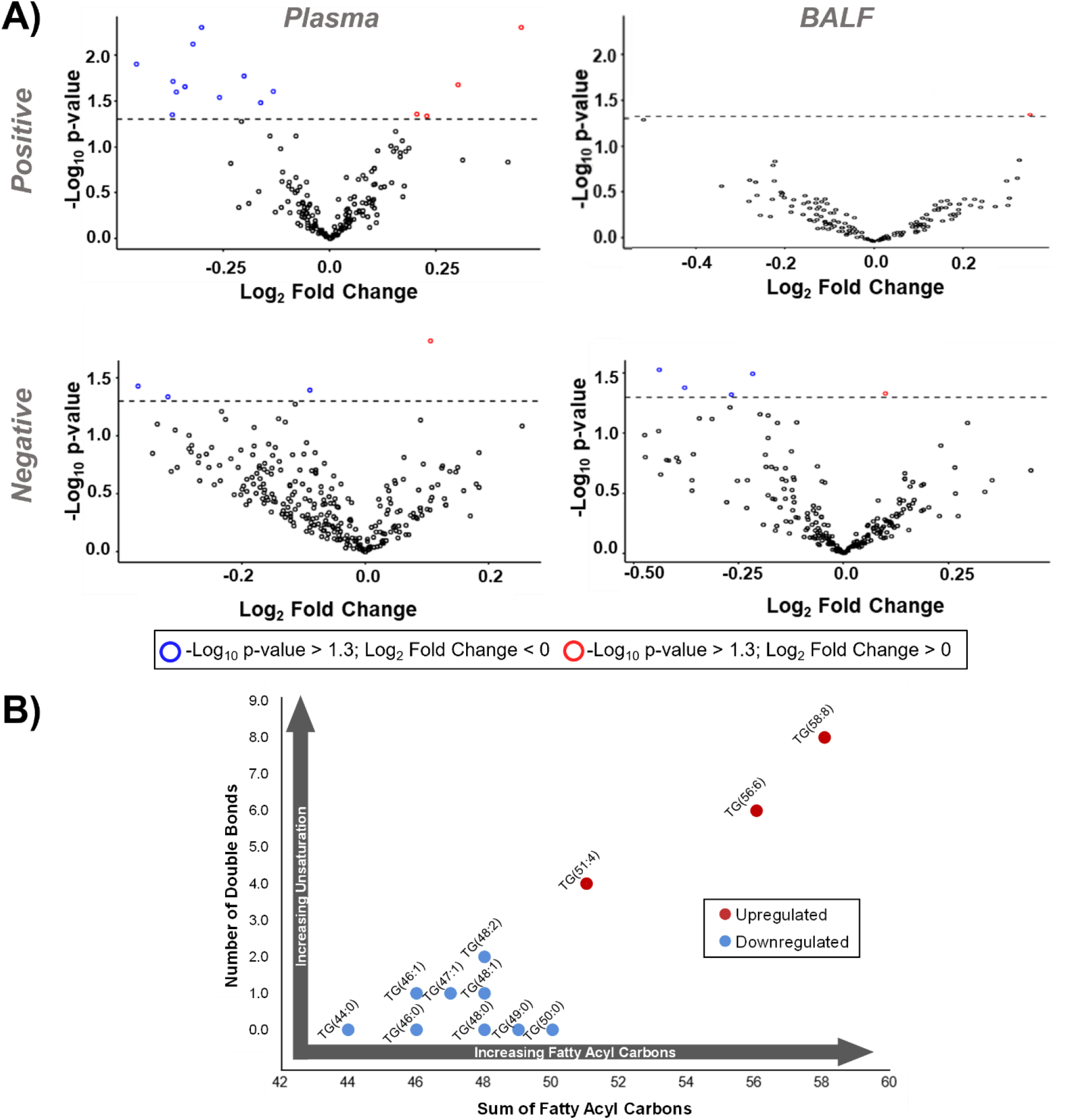
Analysis of lipid significance and associated trends. **A**) Volcano plots of the identified lipids where red and blue outlines indicate up- and downregulated lipids, respectively, in subjects whose smoke inhalation injuries led to mortality. **B**) Association of subject mortality and the fatty acyl composition of statistically significant TGs. When the number of double bonds is plotted against the sum of fatty acyl carbons, a trend emerges in plasma where TGs with fewer carbons and double bonds are downregulated and those with longer, more unsaturated fatty acyls are upregulated in subjects whose injuries led to mortality.

Upon evaluation of the 19 statistically significant lipids in plasma, 5 were found to be upregulated and 14 were downregulated in subjects whose injuries led to mortality (**Table S3**). Dysregulation was specifically observed in plasma TGs, PCs, and PEs, as well as a single DG and PA. Analysis of the lipids showed a clear trend among the dysregulated TGs, as illustrated in **Figure 6B**. Here, TGs with 44-50 fatty acyl carbons and 0-2 degrees of unsaturation were found to be downregulated in subjects whose injuries led to mortality, while TGs with higher numbers of carbons and double bonds were upregulated. This positive correlation between mortality from smoke inhalation injury and both the number of double bonds and carbons of the fatty acyl chains of TGs warrants further investigation as it has also been noted in previous studies.^40, 41^ While the studies noted that the exact pathophysiology and of this complex response is unknown, they have linked this dysregulation to adipose tissue metabolism as burn patients have altered fat metabolism characterized by increased peripheral lipolysis, inadequate hepatic beta-oxidation, and futile cycling of energy substrates including circulating free fatty acids and triglycerides.^42, 43^ In the previous studies, all TGs, however were lumped together and specific species information is not illustrated. Thus, since this TG trend is so pronounced in this study and previous burn analyses, the panel of 12 changing TGs could have prognostic and diagnostic implications for treatment paths of patients.

## CONCLUSIONS

In this manuscript, we developed sample-specific multidimensional lipid libraries using Skyline which were applied to rapidly process and confidently annotate plasma and BALF LC-IMS-CID-MS lipidomic data for the investigation of lipidomic dysregulation related to smoke inhalation injury. The libraries were built from experimental data and stringently validated based on a set of confidence criteria. The two libraries together provided a total of 542 unique lipids, 516 of which were present in the plasma library. Therefore, the plasma library was further developed into a publicly available database which included a Skyline spectral library and iRT calculator. This database leverages many adapted Skyline features such as small molecule spectral libraries, drift time filtering, iRT retention time prediction, analysis of multiple adducts, and neutral loss fragments. In comparison to previous studies of NIST SRM 1950, this workflow when coupled with an LC-IMS-CID-MS platform gave hundreds of confident annotations using a single sample extraction and analysis platform. Upon application to plasma and BALF from 21 patients with smoke inhalation injuries, 25 total lipids were found to be statistically significant, however 19 of these lipids were found in plasma. Based on these findings, plasma appears to be a more robust sample than BALF for prognostic applications as it is much more common, easier to obtain and gave more identified and statistically significant lipids. Of the 19 statistically significant lipids in plasma, a unique trend was observed in the fatty acyl composition of the TGs which correlated to patient mortality. Specifically, TGs with few carbons and less unsaturation were found to decrease in patients whose injuries lead to morality while longer chain and more double bonds increased. While this study was limited by a small number of subject-matched samples, it is the first step in a greater understanding of lipid dysregulation following smoke inhalation and in identifying molecular signatures of injury severity. Furthermore, after additional validation the 12 statistically significant TGs could serve as a biomarker panel for predictive patient prognosis.

## SUPPORTING INFORMATION

LC gradient method; CID ramp method; NIST SRM comparison; spectral library match; iRT reference lipids; co-elution of multiple adducts; BALF and plasma significant lipids (PDF)

Plasma and BALF transition lists (xlsx)

Lipid abundances for statistical analysis (xlsx)

## DATA AVAILABILITY

The Skyline plasma lipid database is publicly available at https://panoramaweb.org/baker-lipid-ims.url. The Skyline files for the NIST SRM and smoke inhalation (plasma and BALF) applications are available at https://panoramaweb.org/lipid-library-applications.url.

## NOTES

The manuscript authors report no competing interests.

## ACKNOWLEDGEMENTS

All measurements were performed in the Molecular Education, Technology and Research Innovation Center (METRIC) at NC State University.

