## Supporting Information for "Development and Application of Multidimensional Lipid Libraries to Investigate Lipidomic Dysregulation Related to Smoke Inhalation Injury Severity"

### TABLE OF CONTENTS

**Table S1.** LC gradient method

**Table S2.** CID ramp method

**Figure S1.** NIST SRM comparison

**Figure S2.** Spectral library match

**Table S3.** iRT reference lipids

**Figure S3.** Co-elution of multiple adducts

**Table S4.** BALF and plasma significant lipids

**Supplemental file 1.** Plasma and BALF lipid library transition lists (separate file)

**Supplemental file 2.** Lipid abundances for statistical analysis (separate file)

**Table S1:** LC gradient method

| Time (min) | % MPA/% MPB | Flow rate (mL/min) |
|---|---|---|
| Elution Gradient | | |
| 0 | 60/40 | 0.25 |
| 2 | 50/50 | 0.25 |
| 3 | 40/60 | 0.25 |
| 12 | 30/70 | 0.25 |
| 15 | 25/75 | 0.25 |
| 17 | 22/78 | 0.25 |
| 19 | 15/85 | 0.25 |
| 22 | 8/92 | 0.25 |
| 25 | 1/99 | 0.25 |
| 34 | 1/99 | 0.25 |
| Column Wash | | |
| 34.5 | 60/40 | 0.30 |
| 35 | 1/99 | 0.30 |
| 35.5 | 1/99 | 0.30 |
| 36 | 60/40 | 0.35 |
| 37 | 60/40 | 0.30 |
| 38 | 60/40 | 0.25 |

**Table S2:** CID collision energy ramp method

| Drift time | Collision energy |
|---|---|
| 0 | 10 |
| 15 | 14 |
| 19 | 27 |
| 25 | 45 |
| 40 | 52 |
| 50 | 58 |

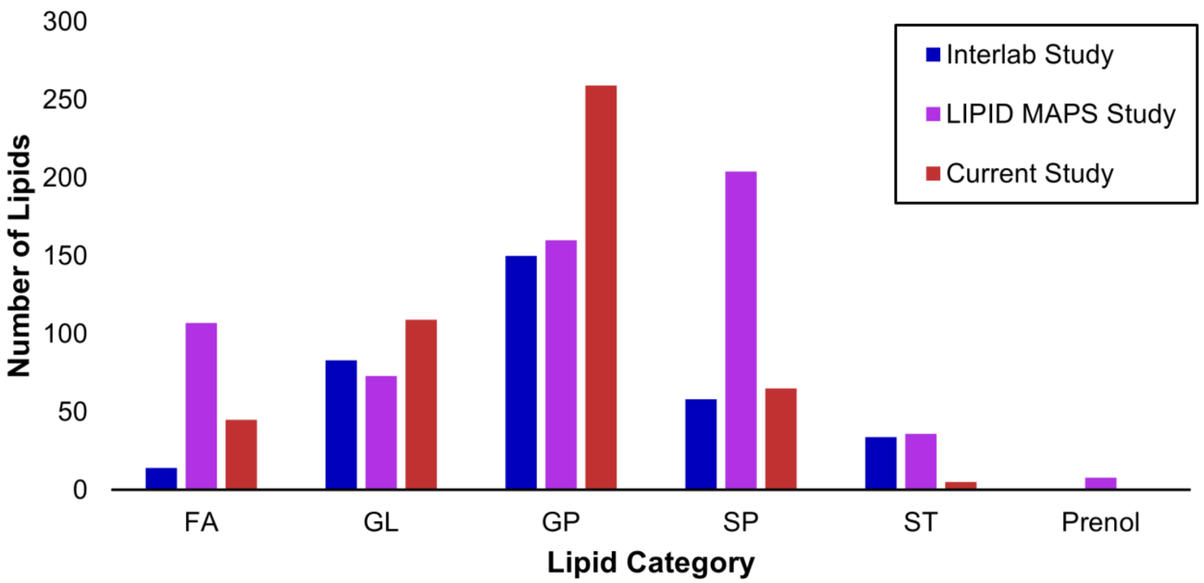


**Figure S1.** Comparison of NIST SRM 1950 results to previous studies. The number of lipids identified per category is plotted where the colors represent the study. Interlab study refers to the interlaboratory comparison from Bowden et al. with only lipids identified in at least 5 of the 31 laboratories, and LIPID MAPS study refers to the LIPID MAPS consortium characterization of SRM 1950 from Quehenberger et al.[32, 33] Sterols were the only category in which our approach identified fewer lipids than both previous studies, likely due to the sterols requiring specializing extractions or gas-phase ionization methods. Our approach gave the highest number of glycerolipids and glycerophospholipids based on isomers which were summed into a single lipid in both previous studies.

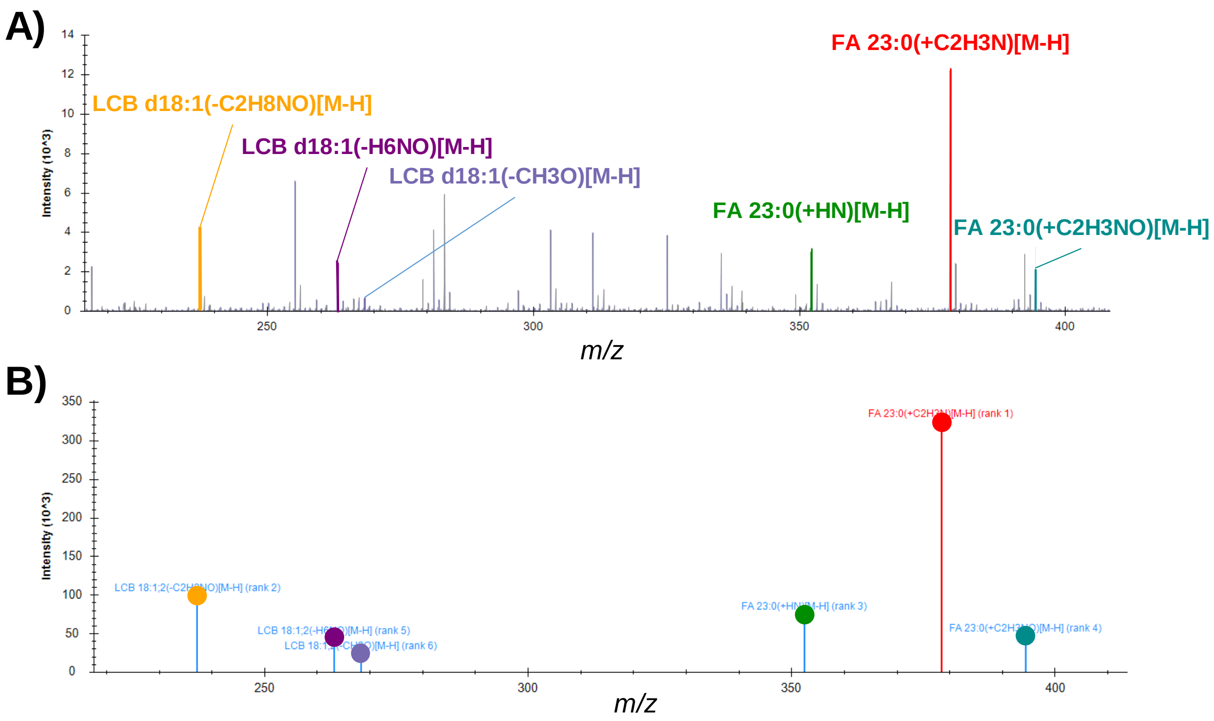


**Figure S2.** Skyline small molecule spectral library match example. **A**) Experimental and **B**) spectral library CID spectra for the lipid Cer(d18:1/23:0). The spectral library was created from pooled plasma whereas the experimental spectra was collected from commercially available BTLE.

**Table S3**: Endogenous iRT reference lipids

| Positive mode | | Negative mode | |
|---|---|---|---|
| **Lipid** | **iRT value** | **Lipid** | **iRT value** |
| AC(10:0) | 0.00 | FA 12:0 | 0.00 |
| AC(18:2) | 11.56 | FA 22:6 | 7.72 |
| AC(18:1) | 15.32 | FA 14:0 | 7.92 |
| AC(18:0) | 20.62 | FA 20:1 | 28.13 |
| SM(d16:1/14:0) | 40.27 | PI(16:0_22:6) | 48.47 |
| SM(d16:1/18:3) | 49.78 | PI(16:0_20:4) | 50.99 |
| SM(d17:1/16:0) | 54.52 | SM(d16:0/16:1) | 56.01 |
| PC(14:0_16:0) | 58.27 | PI(18:0_22:6) | 59.30 |
| PC(O-16:0/18:3) | 64.39 | PI(18:0_20:4) | 61.61 |
| PC(18:0_22:6) | 65.53 | PE(16:0_22:6) | 68.25 |
| SM(d18:1/18:0) | 67.00 | PE(16:0_20:4) | 70.47 |
| PC(18:0_20:3) | 69.81 | PE(16:0_18:2) | 72.31 |
| PC(P-18:0/16:0) | 70.96 | PE(P-16:0/20:4) | 75.88 |
| DG(18:1_18:2) | 73.84 | PC(18:0_22:6) | 75.98 |
| DG(18:1/18:1) | 77.67 | PC(16:0_18:1) | 77.43 |
| TG(60:13) | 87.17 | PE(P-18:0/22:6) | 81.57 |
| TG(16:1_16:1_18:2) | 91.05 | PE(P-18:0/20:4) | 83.50 |
| TG(54:4) | 94.98 | Cer(d18:2/24:1) | 91.96 |
| CE(18:2) | 98.33 | HexCer(d18:1/24:0) | 95.85 |
| TG(18:0_18:1_20:1) | 100.00 | Cer(d18:1/24:0) | 100.00 |

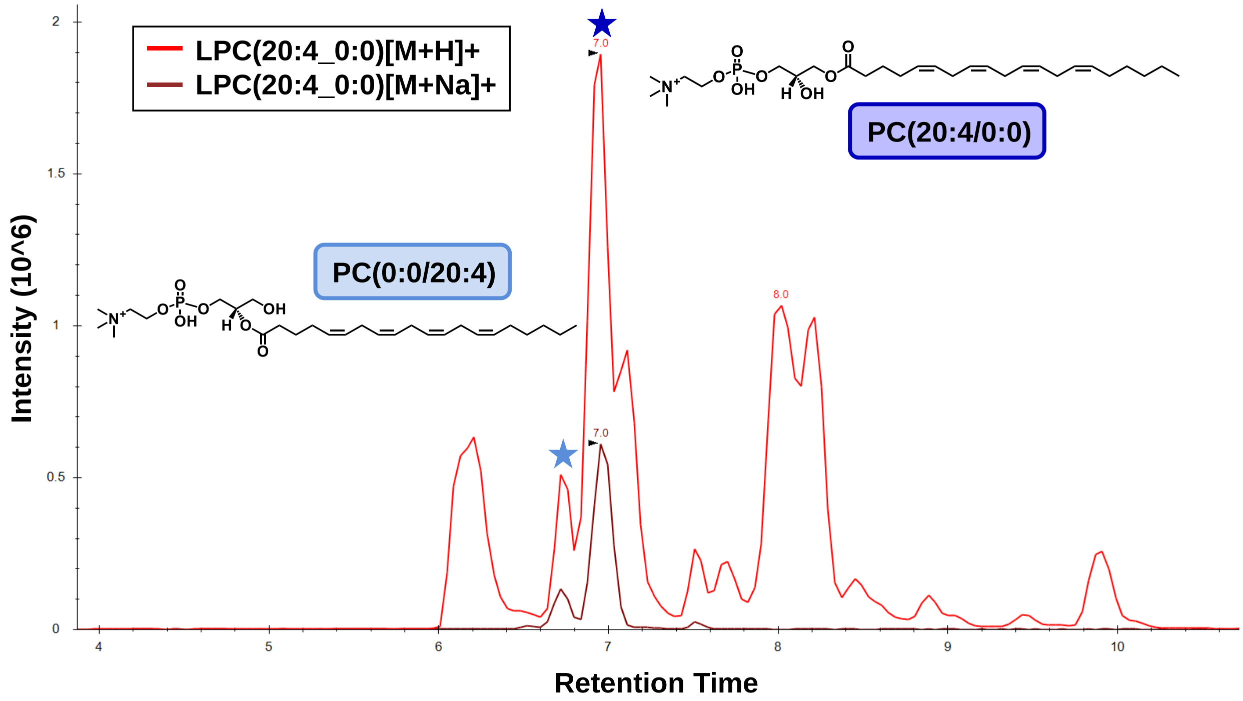


**Figure S3.** Co-elution of multiple adducts aiding in feature annotation. Precursor extracted ion chromatogram (EIC) for LPC 20:4 where two isomeric LC peaks are expected, however the protonated species (red) alone has at least 12 abundant peaks from 6-10 minutes. Addition of the EIC for the sodiated species (brown) gives more confidence that the two features at 6.8 and 7.0 min are LPC(0:0/20:4) and LPC(20:4/0:0), respectively.

**Table S4:** Significant lipids in plasma and BALF

| Sample type | Lipid | Change in subjects whose injuries led to mortality |
|---|---|---|
| BALF | FA 14:0 | Upregulated |
| BALF | AC 14:0 | Upregulated |
| BALF | FA 25:0 | Downregulated |
| BALF | PE(16:0_18:0) | Downregulated |
| BALF | PC(15:0_16:0) | Downregulated |
| BALF | PC(14:0_16:0) | Downregulated |
| Plasma | TG(58:8) | Upregulated |
| Plasma | TG(56:6) | Upregulated |
| Plasma | TG(51:4) | Upregulated |
| Plasma | DG(18:1_18:2) | Upregulated |
| Plasma | PA(16:0_18:1) | Upregulated |
| Plasma | PC(28:0) | Downregulated |
| Plasma | PC(30:1) | Downregulated |
| Plasma | PE(14:0_20:4) | Downregulated |
| Plasma | PE(16:0_18:3) | Downregulated |
| Plasma | PC(18:0_20:2) | Downregulated |
| Plasma | TG(44:0) | Downregulated |
| Plasma | TG(46:0) | Downregulated |
| Plasma | TG(46:1) | Downregulated |
| Plasma | TG(47:1) | Downregulated |
| Plasma | TG(48:0) | Downregulated |
| Plasma | TG(48:1) | Downregulated |
| Plasma | TG(48:2) | Downregulated |
| Plasma | TG(49:0) | Downregulated |
| Plasma | TG(50:0) | Downregulated |
